# The many layers of BOLD. On the contribution of different vascular compartments to laminar fMRI

**DOI:** 10.1101/2021.10.21.465359

**Authors:** Wouter Schellekens, Alex A. Bhogal, Emiel C.A. Roefs, Mario G. Báez-Yáñez, Jeroen C.W. Siero, Natalia Petridou

**Author notes:** **Corresponding author** Name: Wouter Schellekens, Postal address: Q101.132, P.O.Box 85500, 3508 GA, Utrecht, Netherlands.

## Abstract

Ultra-high field functional Magnetic Resonance Imaging (fMRI) offers the spatial resolution to measure neural activity at the scale of cortical layers. Most fMRI studies make use of the Blood-Oxygen-Level Dependent (BOLD) signal, arising from a complex interaction of changes in cerebral blood flow (CBF) and volume (CBV), and venous oxygenation. However, along with cyto- and myeloarchitectural changes across cortical depth, laminar fMRI is confronted with additional confounds related to vascularization differences that exist across cortical depth. In the current study, we quantify how the non-uniform distribution of macro- and micro-vascular compartments, as measured with Gradient-Echo (GE) and Spin-Echo (SE) scan sequences, respectively, affect laminar BOLD fMRI responses following evoked hypercapnic and hyperoxic breathing conditions. We find that both macro- and micro-vascular compartments are capable of comparable theoretical maximum signal intensities, as represented by the M-scaling parameter. However, the capacity for vessel dilation, as reflected by the cerebrovascular reactivity (CVR), is approximately three times larger for the macro-compared to the micro-vasculature at superficial layers. Finally, there is roughly a 35% difference in CBV estimates between the macro- and micro-vascular compartments, although this relative difference is approximately uniform across cortical depth.

## INTRODUCTION

The cortex of the human cerebrum is made up of different layers. The cortical layers can be distinguished on the basis of different neuronal cell types, as well as their connections with other cortical areas or sensory organs^1^. As such, the different cortical layers are hypothesized to account for different sub-processes in brain functioning and human behavior at large^2^. Functional Magnetic Resonance Imaging (fMRI) is one of the most powerful tools for studying brain function in both healthy and diseased individuals non-invasively. Recent advances in ultra-high field MRI (i.e., magnetic field strength ≥ 7 Tesla) now allow for the recording of layer-specific neuronal activity in human populations. The majority of fMRI studies use the Blood-Oxygen-Level Dependent (BOLD) signal to investigate neuronal functions^3^. However, the BOLD signal is an indirect measure of neuronal activity, as it primarily signals differences in the ratio of venous oxy-hemoglobin [Hb] / deoxy-hemoglobin [dHb], affecting T2 and T2* MR-effects. The change in [Hb]/[dHb] ratio following neuronal activity is mainly caused by increases in cerebral blood flow (CBF) and cerebral blood volume (CBV), and the subsequent vessel dilation, in the capillary bed, venules, and larger veins in response to the cerebral metabolic rate of oxygen (CMRO_2_)^4,5^. Despite the indirect representation of neuronal activity, fMRI BOLD is known to correspond well with neuronal electrophysiological recordings (Local Field Potentials (LFP) in particular) in both animals and humans^6,7^. Thus, the differences in CBF/CBV that exist for differently sized venous vascular compartments (i.e., capillaries, venules and veins) do not prevent the utilization of the BOLD signal as an adequate proxy for neuronal activation in conventional fMRI studies. However, laminar fMRI measurements are affected by the vasculature on a whole new level. Because the vascular architecture changes across cortical depth, a confounding correlation arises between different vascular compartments and the different cortical layers^8,9^. This is particularly problematic for the fMRI BOLD scan acquisition sequence that is most commonly used due to its superior sensitivity: Gradient-Echo (GE) BOLD. GE BOLD is sensitive to all venous vascular compartments, but the signal scales with vessel diameter^10,11^. GE BOLD, therefore, is disproportionally sensitive to larger (draining) veins, which predominantly reside near the cortical pial surface. This relative hypersensitivity to the macro-vasculature leads to a relative increase in raw BOLD signals measured at superficial layers, while simultaneously suffering from a decrease in neuronal specificity, as the largest veins pool blood from extended regions of cortex. Therefore, even the scaling of the raw BOLD signal to e.g., percent signal change, cannot prevent or neutralize the fact that neuronal populations of different sizes, represented through different vascular compartments, contribute to the BOLD signal differently across cortical depth^12^. The field of laminar fMRI is currently lacking an adequate quantification of the effect of different vascular compartments on the BOLD signal at the laminar level. Here, we address this topic by conducting a series of measurements in which we record from micro- and macro-vascular compartments across cortical depth, while applying respiratory stimuli to characterize the confounding correlation between differently sized vascular compartments and the BOLD signal.

To characterize the influence of different vascular compartments on the fMRI BOLD signal we capitalize on the increased BOLD Contrast-to-Noise Ratio (CNR) afforded by a 7 Tesla MR system along with boosted sensitivity obtained using a high-density surface receive array^13^ to acquire high spatiotemporal resolution images capable of distinguishing cortical layers. Using a computer-controlled gas delivery system, we manipulate CBF/CBV by increasing the arterial pressure of CO_2_, a potent vasodilator, in a controlled manner^14–16^. The increased CBF/CBV decreases the relative venous deoxy-hemoglobin content, which leads to a BOLD signal increase. To modulate oxygen saturation in the absence of vascular responses, we also apply a hyperoxic stimulus. Increasing the inhaled concentration of O_2_ causes a relative increase in the venous concentration of oxy-hemoglobin, which subsequently results in a BOLD signal increase. Beside the estimation of the BOLD signal change as a result of vasoactive stimulation, the hypercapnia and hyperoxia breathing challenges can be wielded to estimate changes in cerebral vascular reactivity (CVR), which represents the capacity for vessel dilation^17–19^, the M-scaling parameter reflective of the theoretical maximal signal change^20,21^, as well as the change in CBV during separate levels of hypercapnia^15,22^. With these parameters we can quantify to what extent the amplitude of the BOLD response is caused by a vessel’s capacity for dilation (CVR), the maximum venous oxygen content (M-scaling), or the relative CBV increase.

In the current study, we investigate the effect of vasoactive stimuli on laminar fMRI BOLD signals originating from different vascular compartments across cortical depth. Where the GE BOLD signal is weighted towards the macro-vasculature, the Spin-Echo (SE) BOLD signal is generally believed to reflect the micro-vasculature (i.e., mostly capillaries) at high field strengths^23–25^. Unlike the macro-vasculature, the micro-vasculature is uniformly distributed across cortical depth, and is not believed to be capable of vessel dilation in a similar fashion as larger veins^8,9,26^. We utilize GE and SE scan sequences at approximately laminar spatial resolution as measures of macro- and micro-vascular compartments, respectively. Hypercapnia and hyperoxia conditions are realized during scanning to characterize the effects of vasoactive stimuli on different vascular compartments. We expect a percent signal BOLD increase (i.e., %ΔBOLD) for all vascular compartments (i.e., GE & SE), during both hypercapnic and hyperoxic breathing conditions (i.e., increased levels of CO_2_ and O_2_) across cortical depth. However, the %ΔBOLD as well as CVR, M-scaling, and ΔCBV sampled from the macro-vasculature are hypothesized to increase from deep to superficial layer estimates, but not for the micro-vasculature. Finally, macro-/micro-vasculature ratios for CVR, M-scaling, and ΔCBV are calculated, describing the effective relative contribution of the vascular compartments to laminar BOLD fMRI.

## METHODS

### Participants

Eleven healthy volunteers (N = 11, age range 18-42y, mean age = 24.3y, Female = 8) participated in this study after giving written informed consent. All participants declared that they did not experience breathing difficulties under normal conditions, and had not been diagnosed with (cerebro)vascular-related illnesses. The experimental protocol was approved by the local ethics committee of the University Medical Center Utrecht (UMCU) in accordance with the Declaration of Helsinki (2013), and the Dutch Medical Research Involving Human Subjects Act.

### Breathing protocol

During the acquisition of the functional BOLD time-series (see details below), we administered specific breathable gas mixtures to the participants. Hypercapnia and hyperoxia conditions were achieved by increasing the CO_2_ and O_2_ gas concentrations, respectively. Postapneic End-tidal (Pet)CO_2_ and PetO_2_ pressure values were targeted using a computer-controlled gas blender and sequential gas delivery system. (3^rd^ generation RespirAct™, Thornhill Research Inc, Toronto, Canada). A 697s breathing task was performed consisting of the following 4 parts: (1) 200s baseline period with subject-specific targeted PetCO_2_ values. (2) 120s hypercapnia period of +3 or +5 mmHg PetCO_2_ increase. (3) 120s hypercapnia period of +8 or +10 mmHg PetCO_2_ increase. (4) 120s hyperoxia period of +350 mmHg PetO_2_ increase (Figure 1). The breathing task was performed twice by all participants: once for each scan acquisition sequence (i.e., GE & SE), during which the hypercapnia conditions consisted of a +5 mmHg PetCO_2_ increase followed by a +10 mmHG PetCO_2_ increase. The experiment was repeated in 7 participants with +3 mmHg PetCO_2_ and +8 mmHg PetCO_2_ hypercapnia conditions both for GE and SE scan acquisitions. The subject-specific baseline PetCO_2_ calibration was estimated before scanning.

**Figure 1.**
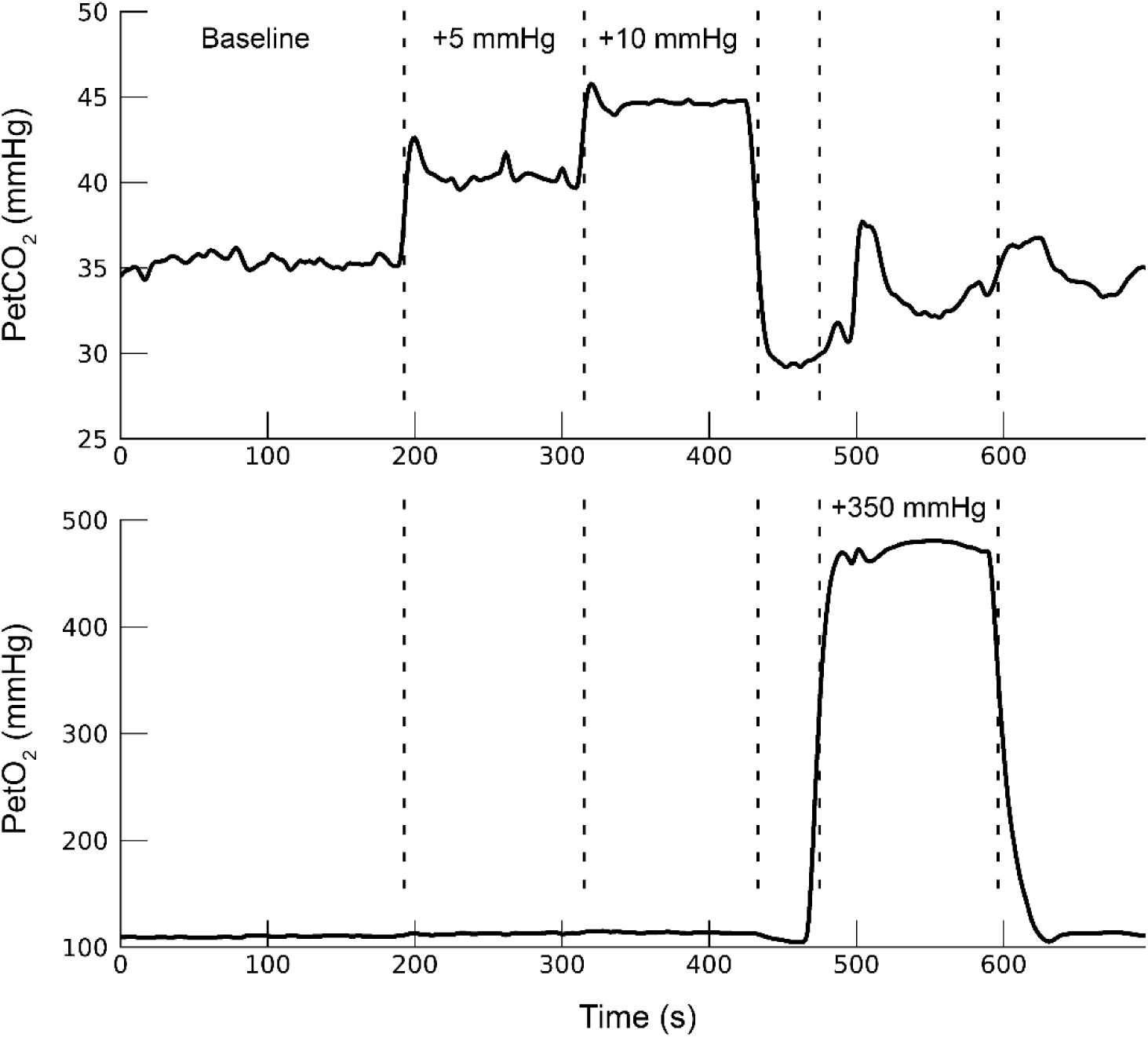
Breathing protocol. For 1 participant (subj08) the hypercapnic (top panel) and hyperoxic (bottom panel) breathing conditions are shown. The dashed lines mark the different parts of the breathing protocol.

### Scan protocol

Scanning was performed at the UMCU on a 7T Philips Achieva scanner (Philips Healthcare, Best, the Netherlands) with two 16-channel high-density surface receive arrays (MRCoils BV, the Netherlands). The anatomical scans consisted of a T1-weighted structural volume: Field Of View (FOV) (ap × fh × rl) = 40 × 159 × 159 mm, which covered the posterior part of the brain (occipital lobe/early visual cortex). The acquired voxel size was: 0.8 × 0.8 × 0.8 mm isotropic, TR/TE = 7.0/2.97ms. T1-weighted volumes at high field strength can experience substantial intensity inhomogeneities. Therefore, a proton density (PD) volume of equal dimension was recorded to correct for these large-scale intensity inhomogeneities. Finally, three T2*-weighted flow-compensated anatomical volumes were acquired with similar coverage: FOV (ap × fh × rl) 40 × 161 × 161 mm and 0.5 × 0.5 × 0.5 mm voxel size, TR/TE = 56/30ms. Both magnitude and phase volumes were reconstructed.

Functional volumes were acquired with two different scan sequences; GE and SE sequences, both using Echo Planar Imaging (EPI). The GE volumes were acquired with SENSE-factor = 3.0, EPI-factor = 31, TR/TE = 850/27ms, flip-angle (FA) = 50°, voxel size = 1.0 × 1.0 × 1.0 mm, FOV = 7 × 128 × 128 mm covering a portion of early visual cortex within the occipital lobe. To increase the Signal-to-Noise Ratio (SNR), the SE volumes were required at a lower spatial resolution: SENSE-factor = 2.0, EPI-factor = 63, TR/TE = 850/50ms, FA = 90°, voxel size = 1.5 × 1.5 × 1.5 mm, FOV = 7.5 × 190 × 190 mm. During a single MRI-session, a maximum of four fMRI time series (i.e., 2 × GE and 2× SE) were recorded, during which the different breathing protocols were applied. Each time series consisted of 820 volumes (duration = 697s per time series). For both GE and SE sequences 5 volumes with reversed phase encoding (i.e., right-left) were recorded to correct for geometric distortions. During all acquisitions, respiration was measured with a respiratory belt around the chest, and blood pulsation with a peripheral pulse unit (PPU). The respiration and PPU measurements were used to calculate the Respiration Volume per Time (RVT) and beats per minute (BPM)^27^.

### Preprocessing

The T1-weighted volume was divided by the PD volume to correct for large-scale intensity inhomogeneities^28^. Afterwards, the T1 weighted volume was resampled to a resolution of 0.2 mm^3^ isotropic voxel size to estimate cortical layers at high spatial resolution (Figure 2). The cortex was divided into 20 equivolumetric laminae using the LayNii software package^29^. Here, the word ’laminae’ is used rather than ’layers’ to emphasize that these laminae do not represent architectonic layers distinguishable with histology, but reflect a measure of cortical depth.

**Figure 2.**
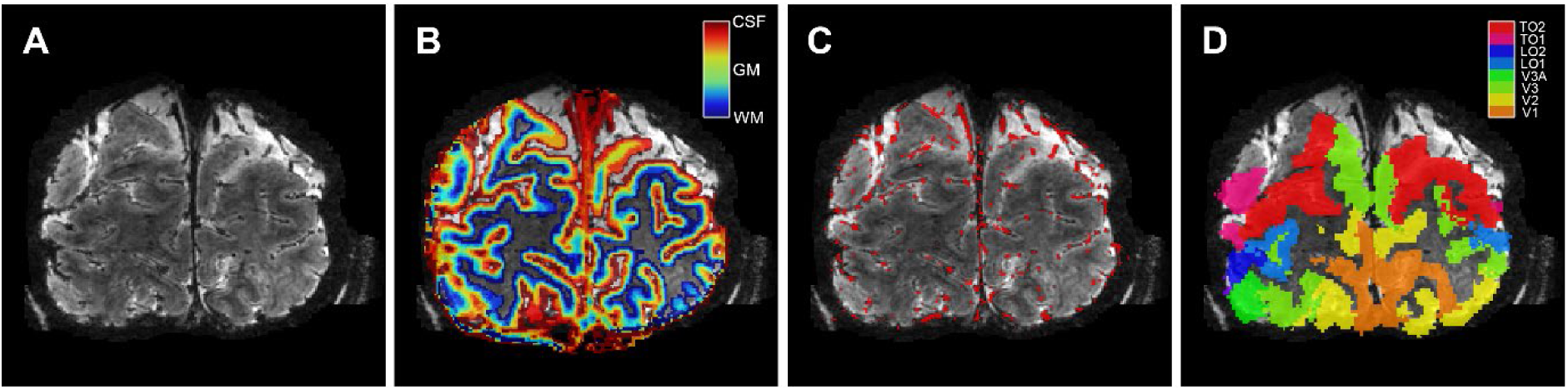
Volumetric maps. For 1 participant (subj09) the T2*-weighted anatomical volume (A), the layer segmentation (B), the pial vein estimation (C), and the visual ROIs (D) are shown. Of the visual ROIs, only V1, V2, and V3 were included.

The three T2*-weighted volumes were first realigned and averaged to increase signal-to-noise. The mean T2*-weighted volume was used to segment large veins, which have a near-zero intensity due to the low T2 of blood and, therefore, appear black within the volume (Figure 2A). The vein segmentation was performed twice with different software packages that produced complementary results. First, large veins were estimated on the magnitude volume with Braincharter^30^. Second, large veins were estimated again with Nighres^31^ on a quantitative susceptibility map (QSM). The QSM was reconstructed by Laplacian-based unwrapping and SHARP background filtering of the phase volume^32,33^, and subsequently an iterative rapid two-step dipole inversion method^34^. Both methods were combined to obtain the pial vein estimation volume (Figure 2C).

A Region-of-Interest (ROI) approach was adopted, consisting of the primary visual cortex (V1) and extra-striate areas V2 and V3, as these areas are believed to have a comparable vascularization^35^. Estimates of early visual cortical areas V1, V2, and V3 were constructed using a whole-brain 3 Tesla T1-weighted volume that was available for the participants. A white and grey matter cortical surface was estimated on the 3T T1-weighted volume with Freesurfer (https://surfer.nmr.mgh.harvard.edu). The cortical surface reconstructions were then used to generate a surface-based visual area maps using the anatomically defined Benson atlas of visual areas with Neuropythy (https://github.com/noahbenson/neuropythy)^36^. The visual area maps were projected back to volumetric space, and through a co-registration of 3T and 7T T1-weighted volumes, transformed to 7T T1-weighted space (Figure 2D).

All functional volumes were corrected for rigid body head motion with AFNI’s 3dvolreg. The EPI phase-encoding induced geometric distortions were corrected using AFNI’s 3dQwarp. This EPI distortion correction and the motion correction were simultaneously applied in a single interpolation step using 3dNwarpApply to generate motion-corrected undistorted functional time series^37,38^. An affine registration was then performed between the mean volume of the functional time-series and the T1 anatomical volume using antsRegistration (http://stnava.github.io/ANTs/)^39^. The inverse of this transformation matrix was used to transform the previously acquired laminae volume, pial vein volume, and early visual cortex volume to the origin and dimensions of the functional volumes using a nearest neighbor interpolation. Lastly, we spatially smoothed the data per cortical depth level (i.e. deep, middle, and superficial levels) and visual area using a Gaussian kernel with a standard deviation of 1mm. This procedure prevents the blurring of voxel data between different cortical depth levels or visual areas. The functional time series were then high-pass filtered using a discrete cosine transform filtering with a cut-off at 0.01 Hz and re-scaled to percent signal change.

### FMRI data analysis

Estimates of the change in percent BOLD signal for each of the hypercapnia and hyperoxia levels were calculated using a General Linear Model (GLM). The GLM regressors consisted of a binary time series for each available breathing condition, and a set of nuisance regressors consisting of 6 rigid-body head motion parameters and the RVT and BPM. We use binary regressors for the breathing conditions (i.e., value of 1 during the respective condition, 0 otherwise) in order to split up the connected hypercapnia conditions, and obtain regression coefficients for both. A second benefit of the binary gas condition regressors is that they are not affected by transient signal changes (e.g. caused by movements), but rather fit the average plateau of the 2 minute hypercapnia and hyperoxia conditions. The regression coefficients per breathing condition serve as %ΔBOLD for each voxel. The %ΔBOLD values for the hypercapnia conditions (i.e., %ΔBOLD_hc_) were then used to calculate the Cerebral Vascular Reactivity (CVR). For each voxel within participants, a linear regression between the obtained PetCO_2_ increase (i.e., ΔPetCO_2_) and %ΔBOLD_hc_ was performed. The slope of the linear regression within each voxel represents the CVR for that voxel.

To estimate the relative change in CBV, we first used the %ΔBOLD values from the hyperoxia condition (%ΔBOLD_ho_) to estimate the M-scaling factor using the hyperoxia-calibrated BOLD model (including the usage of literature standard values) from Chiarelli et al.^40^:

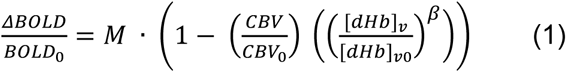

Where M is a scaling parameter that represents the theoretical maximum signal change. The subscript *“0”* refers to baseline conditions and the subscript *“v”* refers to venous properties. The change in CBV relates to the change in CBF following the venous coupling exponent *α* = 0.2^41^:

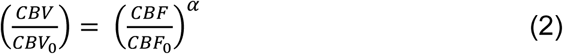

Because hyperoxia is generally believed to have a negligible effect on the change in CBF, equation (1) under hyperoxia conditions can be simplified to:

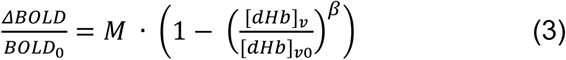

This means that the change in BOLD signal under hyperoxia conditions is caused by the change in venous de-oxyhemoglobin concentration. The change in [dHb]_v_ can be estimated through standard formulas of oxygen transportation in the blood and by assuming a baseline oxygen extraction fraction (OEF). The end-tidal oxygen pressure values can be used to infer the arterial oxygen tension (Pa_O2_). Then, using the Severinghaus equation we can obtain the arterial oxygen saturation (Sa_O2_)^42^:

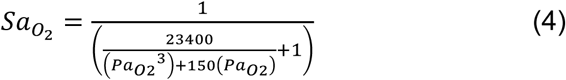

Now, the arterial oxygen content (Ca_O2_) can be estimated by assuming literature standard values for: the O_2_-carrying capacity of hemoglobin (φ: 1.34 ml O_2_ / g_hb_ in humans), the concentration of hemoglobin ([Hb]: 15 g Hb / dl blood), and the solubility coefficient of oxygen in blood (ε: 0.0031 ml O_2_ / (dl blood * mmHg):

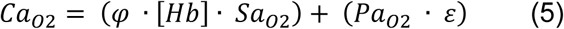

The venous oxygen content (Cv_O2_) depends on the Ca_O2_ and the OEF. We did not measure the OEF, but assume a literature standard value of OEF = 0.30 ^40,43^:

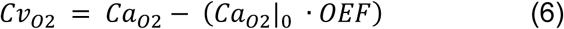

The venous oxygen saturation (Sv_O2_) can be estimated as follows:

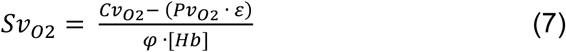

In equation (7), the Pv_O2_ represents the oxygen dissolved in venous plasma and is believed to have a negligible small effect. At this point we can estimate the deoxygenated fraction of [Hb] (F_[dHb]_) from Sv_O2_:

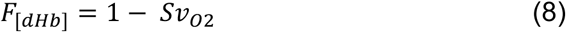

The relative change in F_[dHb]_ during hyperoxia conditions represents the [dHb]_v_/[dHb]_v0_ ratio from equation (3), which means that the M scaling parameter can be estimated as follows:

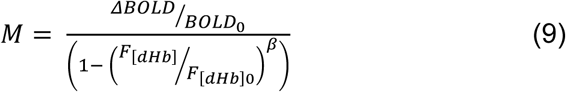

The *“β”* represents the influence of deoxygenated hemoglobin on transverse relaxation, and is estimated at *β* ≈ *1* for 7 Tesla MRI^44,45^.

Now with the estimated M-parameters from the hyperoxia condition, we can estimate the change in venous CBV. The Davis model describes the change in [dHb] as equal to CMRO_2_ and CBF ^20,21^:

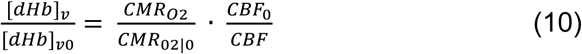

Using equations (2) and (10), we can transform equation (1) to:

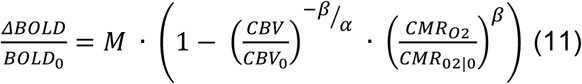

Hypercapnia conditions cause a small metabolic decrease, and was previously estimated to be approximately a 15% decrease for 90% CBF increase (+22 mmHg CO_2_). The effect is believed to scale linearly with CBF increase (and therefore with CO_2_ inspiration), which is why we adopt the following values for 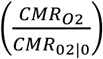 during +3 mmHg, +5 mmHg, +8 mmHg, and +10 mmHg petCO_2_: [0.97; 0.95; 0.92; 0.90], respectively^46^. Now with the estimated M parameter from the hyperoxia condition, we can estimate ΔCBV as follows:

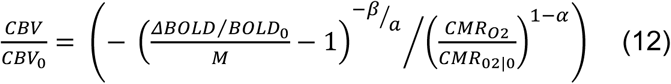

### Statistical analysis

A general linear model (GLM) was constructed that consisted of gas challenge regressors and nuisance regressors (i.e., motion & physiology parameters). Hypercapnia and hyperoxia condition t-statistics were calculated on the basis of regression coefficients for the individual gas challenges. Only voxels that responded significantly to the gas-challenges were selected for further analyses (p < 0.05, Holm-Bonferroni corrected). Additional masks were imposed by the ‘laminae mask’ and ‘visual area mask’, which meant that only voxels were selected that were in range of grey matter cortical layers within visual areas V1, V2, and V3.

Separate linear mixed models (LMM) were constructed with “%ΔBOLD_hc_”, “%ΔBOLD_ho_”, “CVR”, “M”, “ΔCBV” as dependent variables, and with the participants as a random-effects grouping factor. The usage of an LMM analysis allows for the inclusion of each voxel as a separate observation for each of the metrics. Additionally, the model is capable of handling missing values for +3 mmHg and +8 mmHg PetCO_2_ hypercapnia levels in 4 participants, which means that all conditions and measurements of all participants could be included. Each LMM had the following ‘fixed effects’ variables: scan sequence (i.e., GE, SE); and cortical depth (i.e., scaled laminae estimate). The LMM for %ΔBOLD_hc_ and ΔCBV, additionally, have a PetCO_2_ fixed effect variable (i.e., scaled variable of measured PetCO_2_). Random slopes were estimated across the participant random effect. The LMMs were fitted using the restricted maximum likelihood (REML) approach, and the degrees of freedom were calculated using the Satterthwaite model. The statistical test were performed using *JASP* (V.0.15, www.jasp-stats.org).

## Results

### Percent signal change

We observed an average increase in percent signal change following the hypercapnia conditions (F_(1,9.9)_ = 54.17, p < .001). The %ΔBOLD_hc_ differed significantly per scan sequence (F_(1,9.9)_ = 5.40, p = .043. Figure 3), meaning that the micro- and macro-vasculature on average produced significantly different BOLD signal amplitudes (mean %ΔBOLD_hc_ GE = 3.85, 95% CI = [3.58, 4.12]; mean %ΔBOLD_hc_ SE = 2.62, 95% CI = [2.39, 2.84]). While taking both GE and SE into account, there was no effect of cortical depth on %ΔBOLD_hc_ (F_(1,9.3)_ = 1.27, p = .288). However, there was a strong interaction effect of the signal amplitude during hypercapnia levels with the laminae estimates (F_(1,10.0)_ = 29.06, p < .001), and the additional three-way interaction with scan sequence (F_(1,10.4)_ = 20.32, p = .001). These results indicate that %ΔBOLD_hc_ increases more strongly with increased CO_2_ inspiration at superficial layers than deeper layers. This effect was more prominent for the GE scan sequence as opposed to the SE scan sequence (Figure 4).

**Figure 3.**
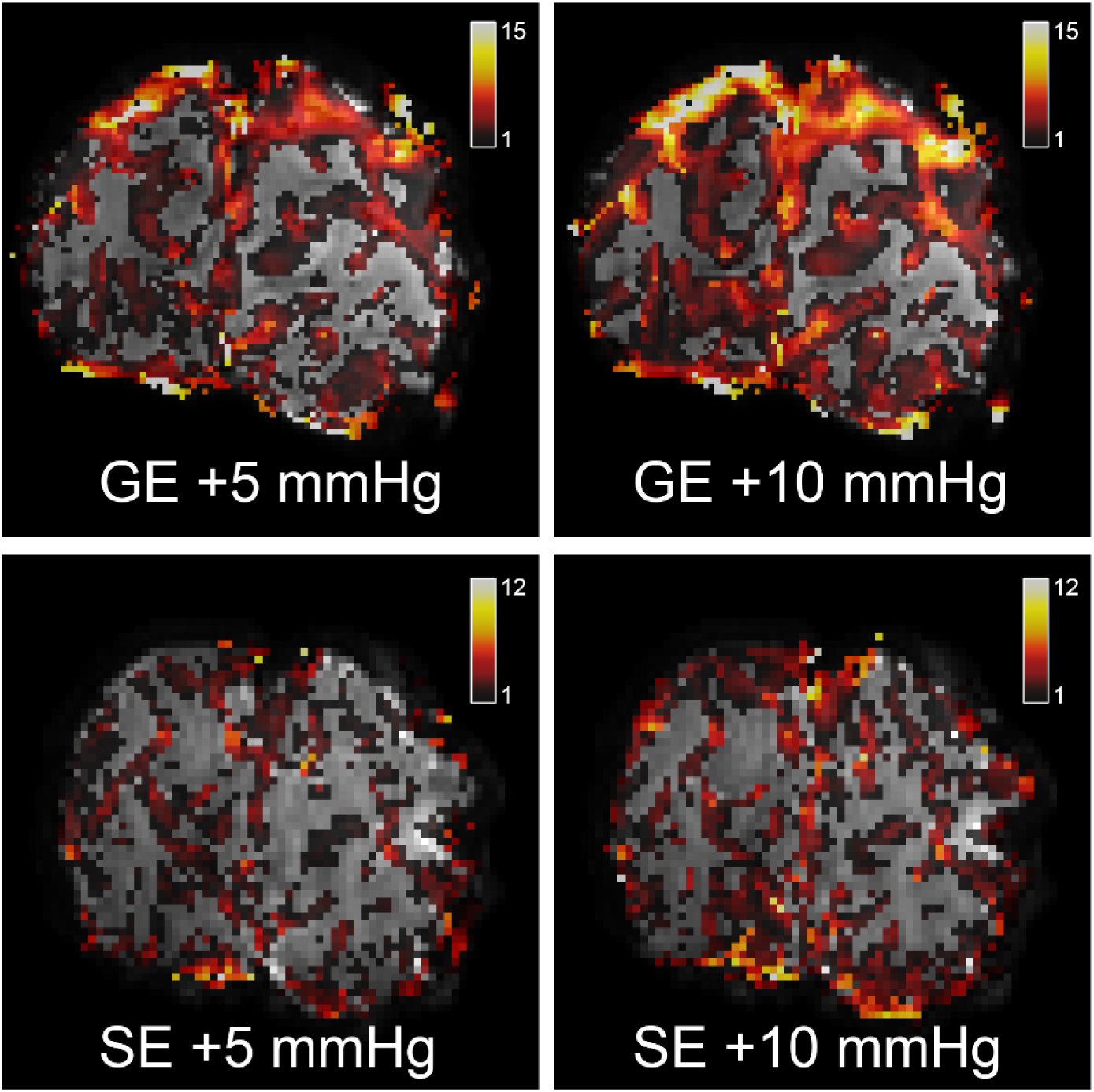
Volumetric hypercapnia BOLD effect. The %ΔBOLDhc is shown for one participant (subj02) for 2 hypercapnia levels: +5 mmHg & +10 mmHg PetCO2 (left-right panels), as measured with GE (top panels) and SE (bottom panels).

**Figure 4.**
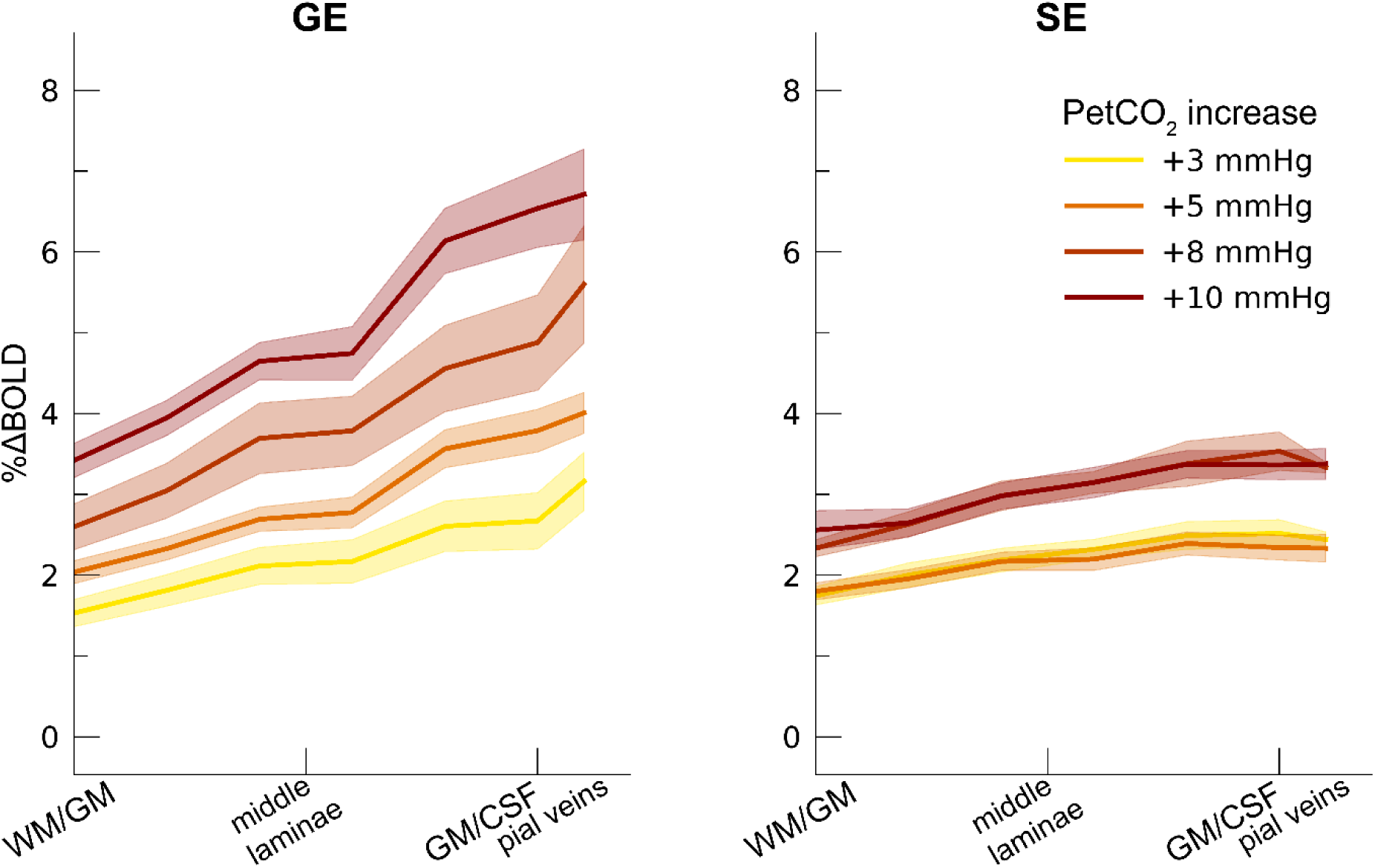
Percent BOLD signal change hypercapnia. The %ΔBOLDhc is shown across cortical depth for the 4 hypercapnia levels (colors) and 2 scan sequences (left/right panels). The shaded area represents the SEM across participants.

The hyperoxia condition also influenced the BOLD signal amplitude %ΔBOLD_ho_ (t_(7.7)_ = 8.97, p < .001; mean %ΔBOLD_ho_ GE = 2.48, 95% CI = [1.89, 3.07]; mean %ΔBOLD_ho_ SE = 2.28, 95% CI = [1.92, 2.64]). In contrast to the hypercapnia conditions, the %ΔBOLD_ho_ increased from deeper to superficial laminae during both GE and SE scan sequences (F_(1,9.7)_ = 68.72, p < .001), without there being a difference observed between scan sequences (F_(1,9.6)_ = 3.90, p = .078), nor an interaction of scan sequence and laminae (F_(1,9.2)_ = 5.01, p = .052). These results signify that both the micro- and macro-vasculature are highly sensitive to the relative increase in venous oxyhemoglobin, having the largest effect near the pial surface (Figure 5).

**Figure 5.**
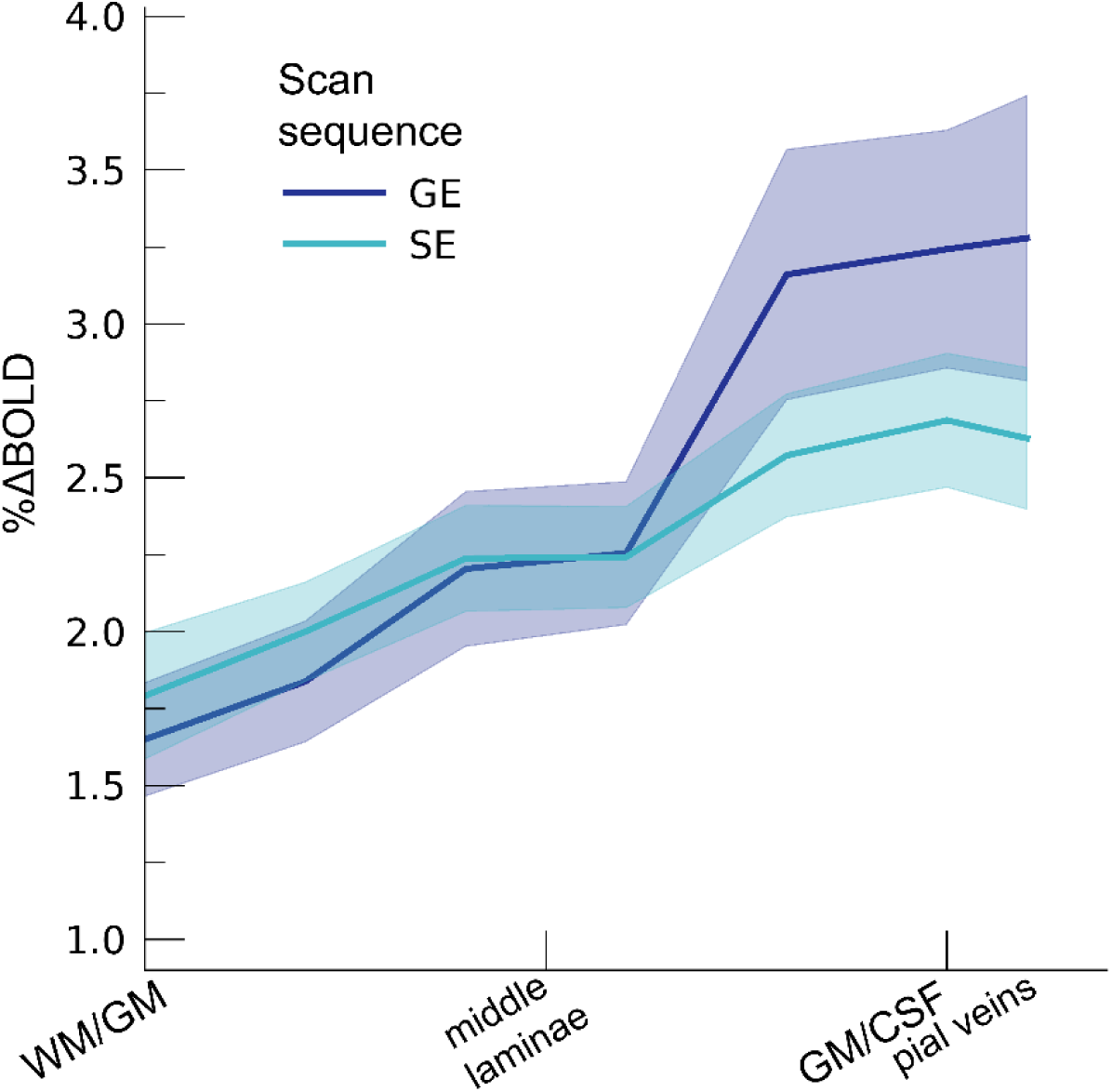
Percent BOLD signal change hyperoxia. The %ΔBOLDho is shown across cortical depth for the 2 scan sequences (colors). The shaded area represents the SEM across participants.

### CVR

The laminae estimates had a significant effect on CVR. Generally, CVR increased towards the superficial layers (F_(1,10.0)_ = 34.16, p < .001). The increase in CVR towards the superficial layers was particularly apparent for the GE scan sequence, as shown by the significant interaction effect between scan sequence and cortical depth factors (F_(1,10.0)_ = 17.02 p = .002). In contrast, the SE sequence did not show a strong increase in CVR from deeper to superficial layers (Figure 6). The deeper cortical layers showed on average a CVR estimate of CVR = 0.39 for GE (95% CI = [0.28, 0.50]) and CVR = 0.18 for SE (95% CI = [0.13, 0.23]), whereas the superficial cortical layers exhibited CVR estimates of CVR = 0.64 for GE (95% CI = [0.50, 0.79]) and CVR = 0.25 for SE (95% CI = [0.20, 0.31]). Thus, the increase in CVR across cortical depth is over a factor of 3 larger for the macro-vasculature than the micro-vasculature.

**Figure 6.**
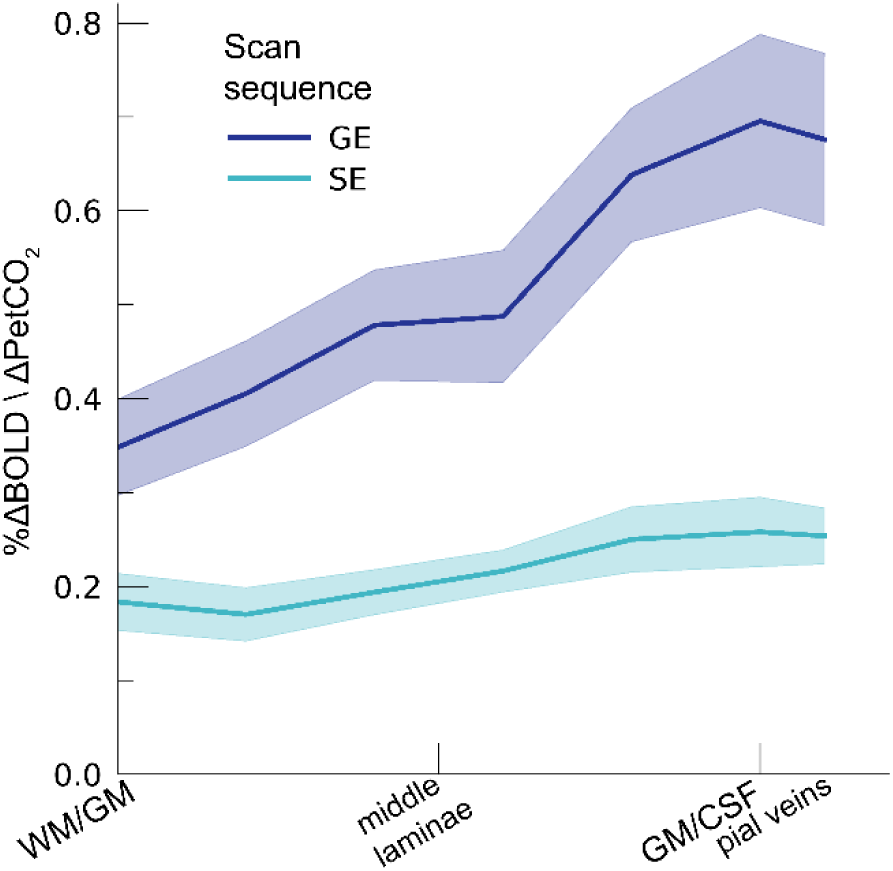
CVR. The CVR is shown across cortical depth for the 2 scan sequences (colors). The shaded area represents the SEM across participants.

### M-scaling

We estimated the M-value based on the %ΔBOLD_ho_ and the PetO_2_ trace during hyperoxia (Figure 7). The M-scaling parameter increased strongly across cortical depth, peaking near the GM/CSF border (F_(1,10.0)_ = 75.79, p < .001). However, a difference between scan sequences was not observed (F_(1,9.9)_ = 0.54, p = .479). M-scaling parameter estimates reveal a substantial maximum signal change capacity for both scan sequences (mean M-scaling GE = 14.28, 95% CI = [11.87, 16.69]; mean M-scaling SE = 12.26, 95% CI = [10.58, 13.94]). We, additionally, observed an interaction effect of scan sequence and cortical depth (F_(1,9.8)_ = 9.88, p = .011). This interaction effect reflects the fact that no difference in M-scaling at deeper cortical laminae between GE and SE sequences was observed (mean M-scaling deeper laminae GE = 10.74, 95% CI = [8.84, 12.65]; SE = 10.38, 95% CI = [8.64, 12.62]; post-hoc z = 0.45, p = .653), while M-scaling was significantly larger at superficial laminae for GE compared to SE (mean M-scaling deeper laminae GE = 17.81, 95% CI = [14.73, 20.88]; SE = 14.14, 95% CI = [12.22, 16.05]; post-hoc z = 2.97, p = .009).

**Figure 7.**
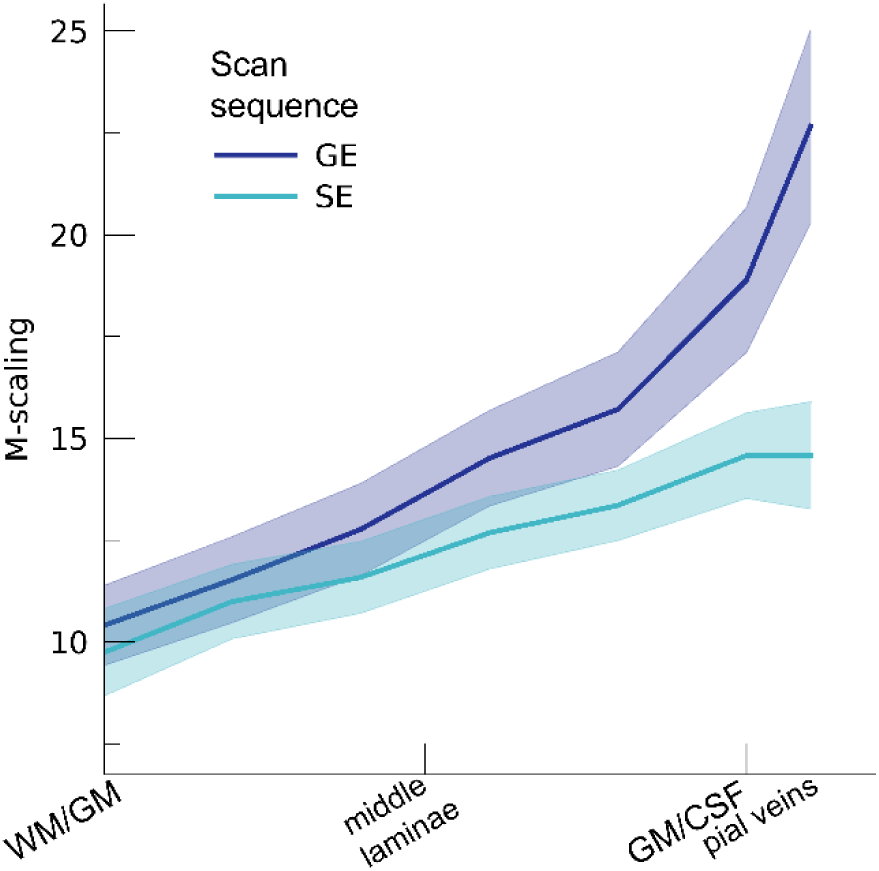
M-scaling. The theoretical maximum signal intensity “M” is shown across cortical depth for the 2 scan sequences (colors). The shaded area represents the SEM across participants.

### ΔCBV

Our ΔCBV estimate shows a significant effect of different hypercapnia levels (F_(1,10.2)_ = 28.63, p < .001), which is representative of an increase in ΔCBV with increased levels of inspired CO_2_ (Figure 8). We, additionally, observed an interaction effect of hypercapnia levels with the scan sequence (F_(1,9.3)_ = 8.61, p = .016) as the ΔCBV increase is approximately 1.35 times larger for GE (mean %ΔCBV = 10.1, 95% CI = [8.1, 12.2]), compared to SE (mean %ΔCBV = 7.4, 95% CI = [6.5, 8.2]). We did not observe a difference in ΔCBV across cortical depth (F_(1,15.5)_ = 0.23, p = .637). Thus, even though the different hypercapnia levels led to a gradual increase in CBV, this relative increase was approximately uniform across cortical depth.

**Figure 8.**
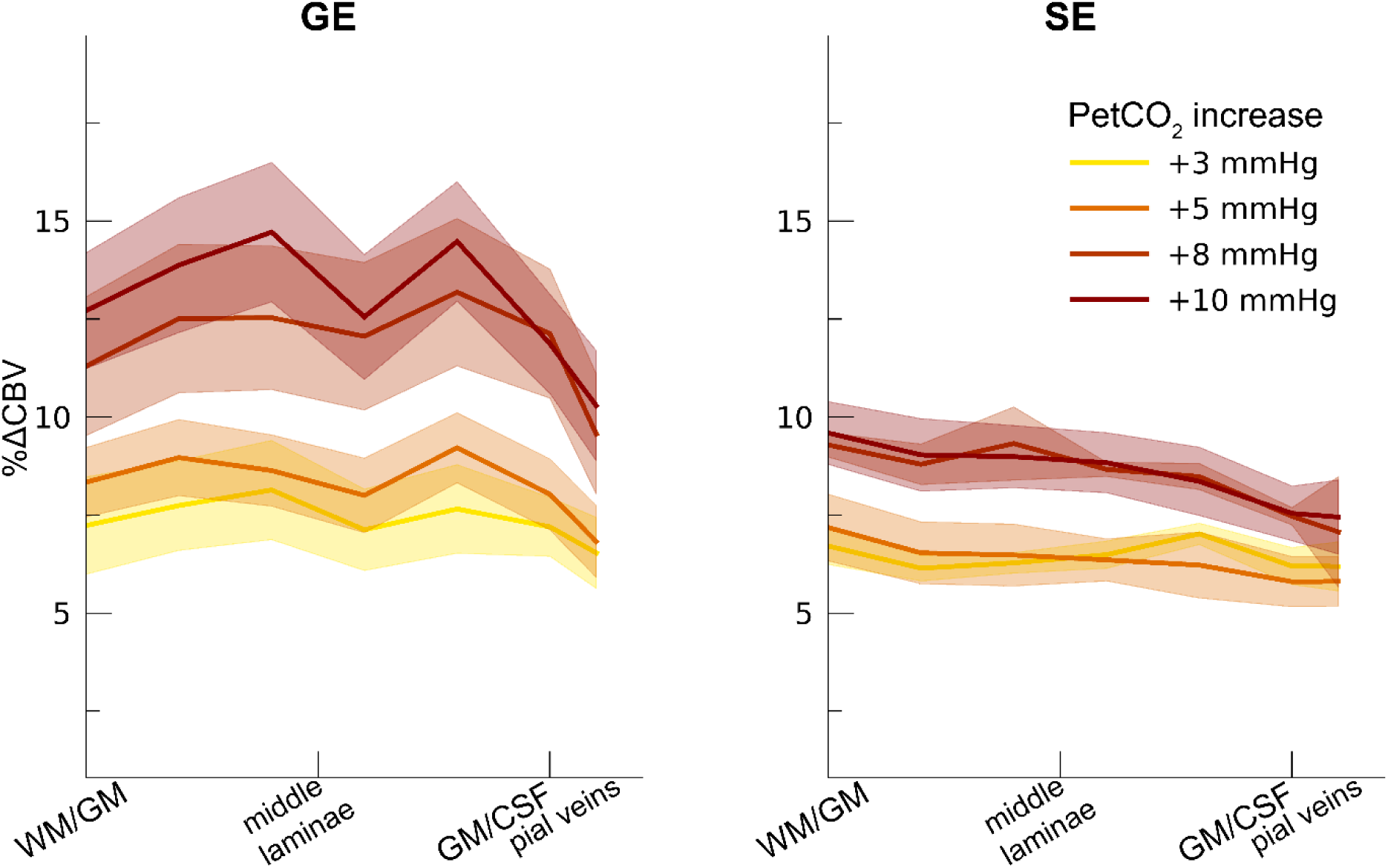
ΔCBV. The ΔCBV in percentages is shown across cortical depth for the 4 hypercapnia levels (colors) and 2 scan sequences (left/right panels). The shaded area represents the SEM across participants.

## Discussion

### General discussion

In the current study, we quantify the different effects that macro- and micro-vascular organization have on laminar fMRI BOLD signal. We find that increasing levels of hypercapnia result in increasing percent signal changes for both the macro- and micro-vasculature. However, the effect of hypercapnia on the BOLD signal is strongly dependent on cortical depth, as well as the different vascular compartments from which the signal originates. This effect is signified by the increasing CVR across cortical depth as sampled from the macro-vasculature. CVR estimates from the micro-vasculature do not show much difference in vessel dilation capacity across cortical depth. The hyperoxia condition also leads to an increase in percent signal change, which together with the PetO_2_ trace allows for an estimation of the maximum theoretical BOLD signal: M-scaling. We find that M-scaling values increase strongly from deeper to more superficial layers. This trend was observed for both the micro- and macro-vasculature, albeit that the trend is significantly steeper for the macro-vasculature. Finally, we observed that increased levels of hypercapnia lead to an increase in ΔCBV, which is significantly more pronounced in the macro-versus the micro-vasculature. We did not observe that the relative change in CBV differs across cortical depth.

### Hyperoxia and the theoretical maximal BOLD signal change

We observe a mean increase in percent BOLD signal change following the hyperoxia condition of +350 mmHg PetO_2_ increase, in line with previous hyperoxia reports^40,47–50^. The increase in PetO_2_ presented here, equates to an air mixture consisting of roughly 60% O_2_, which is considered mild hyperoxia. It has previously been reported that mild cases of hyperoxia have a negligible effect on CBF^47,50,51^. This assumption allows for the estimation of the theoretical maximal percent signal change (M-scaling) per voxel on the basis of the hyperoxia BOLD signal change and the measured PetO_2_ values. We find that both the macro-vasculature as well as the micro-vasculature are capable of generating a large BOLD signal change, purely on the basis of a relative venous oxyhemoglobin increase. Additionally, we find that the theoretical maximal BOLD signal change is lower at the deeper compared to the superficial cortical layers, ranging from 9 – 21 for the macro-vasculature and 9 – 16 for the micro-vasculature in line with previous high-field M estimations^15,45,52^. The M-scaling increase with cortical depth is logically reconcilable with the GE scan sequence, since GE scans are disproportionally sensitive to larger veins near the pial surface, increasing the maximum signal intensity. However, the increase in M-scaling across cortical depth was also seen for the SE scan sequence, albeit with a smaller slope. This either means that the venous oxyhemoglobin increase differs across cortical depth independent from vascular compartment size, or that our SE scan sequence was unintentionally sensitive to vascular compartments larger than the capillary bed only.

### Cerebrovascular reactivity (CVR)

CVR is commonly used to describe vessel dilation properties^17^. Here we show that the macro-vasculature as measured by the GE scan sequence has a greater capacity for vessel dilation than the micro-vasculature as measured by the SE scan sequence. As the largest of the veins reside near the pial surface, we see that CVR increases for the macro-vasculature from deeper towards superficial cortical layers. This effect was not observed for the micro-vasculature. These findings indicate that capillaries and possibly smaller venules have a smaller capacity for dilation. Where similar CVR dilation properties for micro- and macro-vessels are observed for deeper cortical laminae, the macro-vasculature shows on average three times as much capacity for dilation than the micro-vasculature. Since CVR is often interpreted as a proxy for vessel health, high-spatial resolution vessel health measurements based on CVR should correct for the different dilation properties of differently sized vascular compartments across cortical depth. Additionally, the current findings implicate that neural signals as conveyed by the neurovascular coupling from smaller vascular compartments are limited by the maximum dilation capacity of capillaries^26^. The M-scaling parameter, however, indicates that the micro-vasculature is capable of generating a BOLD signal change comparable to the macro-vasculature. The fact that large BOLD signals from smaller vascular compartments are not frequently observed^11,53^, likely stems from the inability of the smallest vessels to dilate in a similar fashion as larger vessels.

### Cerebral Blood Volume (CBV)

Through the measurements of BOLD signal change during hypercapnia levels, the PetCO_2_ trace, and the estimation of the M-scaling parameter, we have been able to estimate the relative change in CBV. A clear increase in CBV is seen for increasing levels of inspired CO_2_, which causes vessels to dilate. A relative increase of 12.5% CBV is seen during the highest hypercapnia level (i.e., +10 mmHg PetCO_2_) for the macro-vasculature. The same hypercapnia level as measured from the micro-vasculature causes on average 8.5% CBV change. Contrary to the other metrics of the current study, we find no significant difference in the CBV change across cortical depth. However, a small dip in ΔCBV around the middle cortical layers can be observed (Figure 5), which has previously been observed with direct ΔCBV measurements, albeit with different experiment conditions^54,55^. Possibly, the absence of clear CBV changes across cortical depth is indicative of a conservation of matter (i.e., blood volume in this case): “what goes in, must come out”. This means that in all cortical layers a comparable relative CBV increase can be expected in early visual cortex following vasoactive stimuli, albeit that the absolute change in CBV likely scales with vessel diameter.

### Limitations

There are several limitations to this study that need to be mentioned. First, we have not been able to obtain all four hypercapnia levels (i.e., +3 mmHg, +5 mmHg, +8 mmHg, and +10 mmHg PetCO_2_) for all participants. All participants have engaged in the +5 mmHg and +10 mmHg PetCO_2_ breathing challenges, meaning that current results of CVR and ΔCBV may be skewed towards these conditions. However, missing values were dealt with by employing LMMs for statistical modeling, thereby including all available observations of the petCO_2_ and estimating random slopes per participant. A second limitation concerns the spatial resolution of the SE scan sequence, which entailed a 1.5 mm isotropic voxel size. This spatial resolution was selected to attain a sufficiently large SNR, but simultaneously increases partial voluming effects that can result in a blurring of cortical laminae and the inclusion of white matter and CSF signals. The last limitation hinges on the assumptions made to estimate the M-scaling parameter and subsequent ΔCBV values. Several of the assumed literature standard values are rarely debated (e.g., the O_2_-carrying capacity of hemoglobin, the concentration of hemoglobin in blood, and the solubility coefficient of oxygen in blood). However, standard values for the *OEF*, change in *CMRO*_*2*_ during hypercapnia, transverse relaxation parameter *β*, and CBF/CBV coupling constant *α* are less well agreed upon. Current M-scaling and ΔCBV results are likely dependent on the assumed parameter values.

### Conclusions

Laminar BOLD fMRI is affected by vascularization differences that exist across cortical depth. In the current study, we reveal that macro- and micro-vascular compartments are capable of generating comparable percent BOLD signal changes across cortical depth. However, the macro-vascular compartments show a threefold capacity for dilation in the superficial cortical layers compared to the micro-vascular compartments. Additionally, the relative change in CBV is 1.35 times larger for the macro-versus micro-vasculature. This finding was unaffected by cortical depth, indicating that the change in CBV is not relatively larger for pail draining veins compared to smaller vessels in early visual cortex.

## Declarations

### Ethics approval

This study was approved by the local medical ethics committee.

### Consent to participate

All participants gave written informed consent prior to inclusion.

### Availability of data, material and code

All data will be accessible through Flywheel.

### Authors’ contribution

Conceptualization: WS, AB, MB, JS, NP

Data acquisition: WS, AB, NP

Analysis: WS, AB, ER

Writing: WS

